# Host circadian clocks do not set the schedule for the within-host replication of malaria parasites

**DOI:** 10.1101/777011

**Authors:** Aidan J. O’Donnell, Kimberley F. Prior, Sarah E. Reece

## Abstract

Circadian clocks coordinate organisms’ activities with daily cycles in their environment. Parasites are subject to daily rhythms in the within-host environment, resulting from clock-control of host behaviours and physiologies, including immune responses. Parasites also exhibit rhythms in within-host activities; the timing of host feeding sets the timing of the within-host replication of malaria parasites. Why host feeding matters to parasites and how coordination with feeding is achieved are unknown. Determining whether parasites coordinate with clock-driven food-related rhythms of their hosts matters because rhythmic replication underpins disease symptoms and fuels transmission.

We find that parasite rhythms became coordinated with the time of day that hosts feed in both wild type and clock-mutant mice, whereas parasite rhythmicity was lost in clock-mutant mice that fed continuously. These patterns occurred regardless of whether infections were initiated with synchronous or with desynchronised parasites.

Malaria parasite rhythms are not driven by canonical clock-controlled host rhythms. Instead, we propose parasites coordinate with a temporally-restricted nutrient that becomes available through host digestion or are influenced by a separate clock-independent host process that directly responds to feeding. Thus, interventions could disrupt parasite rhythms to reduce their fitness, without interference by host clock-controlled-homeostasis.

## INTRODUCTION

Biological rhythms are ubiquitous and allow organisms to maximise fitness by synchronising behaviours, physiologies, and cellular processes with periodicity in their environment. The value of coordinating with daily cycles in, for example, light/dark and temperature in the abiotic environment has long been appreciated, and the importance for parasites of coordinating with rhythms experienced inside hosts and vectors (i.e. the biotic environment), is gaining increasing recognition [1-3]. For example, circadian rhythms in virulence enables the fungal pathogen *Botrytis cinerea* to cope with rhythmic immune defences in plant hosts [4, 5], circadian control of macrophage migration provides incoming *Leishmania major* parasites with more host cells to invade at dusk than dawn [6], and host clocks control the ability of herpes and hepatitis viruses to invade cells and to replicate within them [7, 8]. Parasites that possess their own circadian clocks, such as the fungus *B. cinerea* and protozoan *Trypanosoma brucei* (which causes sleeping sickness), can use their clocks to coordinate with host rhythms [4, 9]. However, it is unclear how coordination with host rhythms is achieved by parasites that are not known to have circadian clocks. For example, malaria (*Plasmodium*) parasites exhibit periodicity in their development during their cycles of asexual replication in red blood cells (the intra-erythrocytic development cycle; IDC). The IDC lasts 24 hours (or multiples of 24 hours, depending on the parasite species) and is characterised by progression through distinct developmental stages at particular times-of-day. For example, the timing of *P. chabaudi’s* IDC transitions are determined by the time-of-day that murine hosts are provided with food [10, 11]. Specifically, parasites remain in early IDC stages when hosts are fasting and complete the IDC at the end of the feeding phase. Why malaria parasites exhibit this schedule is unclear, but coordination with host rhythms and with vector rhythms confers fitness benefits to parasites [12-16].

Explaining how and why malaria parasites complete their IDC according to a schedule is important because cycles of asexual replication are responsible for the severity of malaria symptoms and fuels the production of transmission forms. Furthermore, reports that several mosquito populations are shifting the timing of their blood foraging behaviour to evade insecticide-treated bed nets [17-19], suggest the interaction between parasite and vector rhythms could have complex consequences for disease transmission. Moreover, a mechanistic insight into how the IDC is scheduled may offer insight into developing novel interventions to disrupt parasite replication. Indeed, many antimalarial drugs have increased efficacy towards specific IDC stages [20] and parasites may utilize IDC stage specific dormancy to facilitate survival during antimalarial drug treatment [21]. The extent to which hosts enforce a schedule upon the IDC and to which parasites possess an ability to organise their own IDC with respect to time-of-day cue(s) are open questions [22]. For example, it has been suggested that the IDC schedule is entirely explained by circadian host immune responses killing certain IDC stages at certain times of day [23] or by only allowing access to a nutrient essential to a particular IDC stage at a certain time-of-day [24]. Alternatively, parasites may be able to, at least in part, schedule their IDC to avoid coinciding a vulnerable IDC stage with a dangerous time-of-day, or to maximally exploit a time-limited resource [3]. A foundation for explaining both why the IDC schedule benefits parasites and how it is controlled, requires discovering which of the myriad of host rhythms associated with the time-of-day of feeding are responsible for the IDC schedule.

Here, we use the rodent malaria parasite *P. chabaudi* to test whether the relevant host feeding-associated rhythm is driven by, or independent of, the host’s canonical circadian clock. The canonical mammalian circadian clock operates via a core Transcriptional Translational Feedback Loop (TTFL) involving dimeric proteins that promote the expression of other clock proteins as well as the inhibition of themselves [25]. The feedback and degradation of these proteins forms an oscillator that is entrained via external daily stimuli (Zeitgeber, usually light) to keep the clock precisely tuned to environmental periodicity. TTLF clock-controlled processes include many metabolic pathways relevant to IDC progression. For example, CLOCK and BMAL1 regulate blood glucose levels [26, 27], and melatonin release (which has been suggested to speed up IDC progression [28]). Alternatively, the IDC schedule could simply be driven by the appearance of nutrients/metabolites made available in the blood as a direct consequence of food digestion (i.e. via processes not reliant on clocks). Host-clock-controlled or clock-independent products of digestion could: (i) impact directly on IDC progression by providing essential resources for different IDC stages at different times of day, and/or (ii) provide time-of-day information to the parasite to modulate its rate of development to maximise acquisition of such resources, or (iii) act as a proxy for another important rhythmic factor that the parasite must coordinate with.

We apply time restricted feeding (TRF) protocols to wild type and clock disrupted Per1-Per2 double knockout mice (*Per1/2-*null) and compare the consequences for the IDC of *P. chabaudi* infections initiated with either synchronous or desynchronised parasites. We hypothesise that if a feeding rhythm alone is sufficient to generate an IDC schedule, IDC completion will coincide with feeding in both wild type and *Per1/2*-null TRF mice, and that parasites become (or remain) desynchronised in *Per1/2*-null mice allowed to forage throughout the circadian cycle. In contrast, if feeding rhythms impact on the IDC via host clock-controlled processes, parasites will only be synchronous in wild type mice, regardless of whether initiated with synchronous or desynchronised parasites. In addition to examining the impact of the host clock on the IDC, we also test whether infection of clock-disrupted mice has fitness consequences for both parasites and hosts. The mammalian clock controls many aspects of immunity [29], including the ability of leukocytes to migrate to the tissues [30] and the ability of macrophages to release cytokines [31], and rodents without functioning *Per2* lack IFN-y mRNA cycling in the spleen (a key organ for malaria parasite clearance) and have decreased levels of pro-inflammatory cytokines in blood serum [32]. Thus, we predicted that parasites will achieve higher densities, and hosts experience more severe disease, in *Per1/2*-null compared to wild type mice.

## METHODOLOGY

To investigate whether a functional TTLF circadian clock is required for the IDC of malaria parasites to follow a schedule we performed two experiments. First, we initiated infections with desynchronised parasites to test whether a feeding rhythm alone is sufficient to restore synchrony and timing in the IDC (Figure 1a). Second, we tested whether the loss of rhythmic feeding causes desynchronization in the IDC of infections initiated with synchronized parasites (Figure 1b).

**Figure 1.**
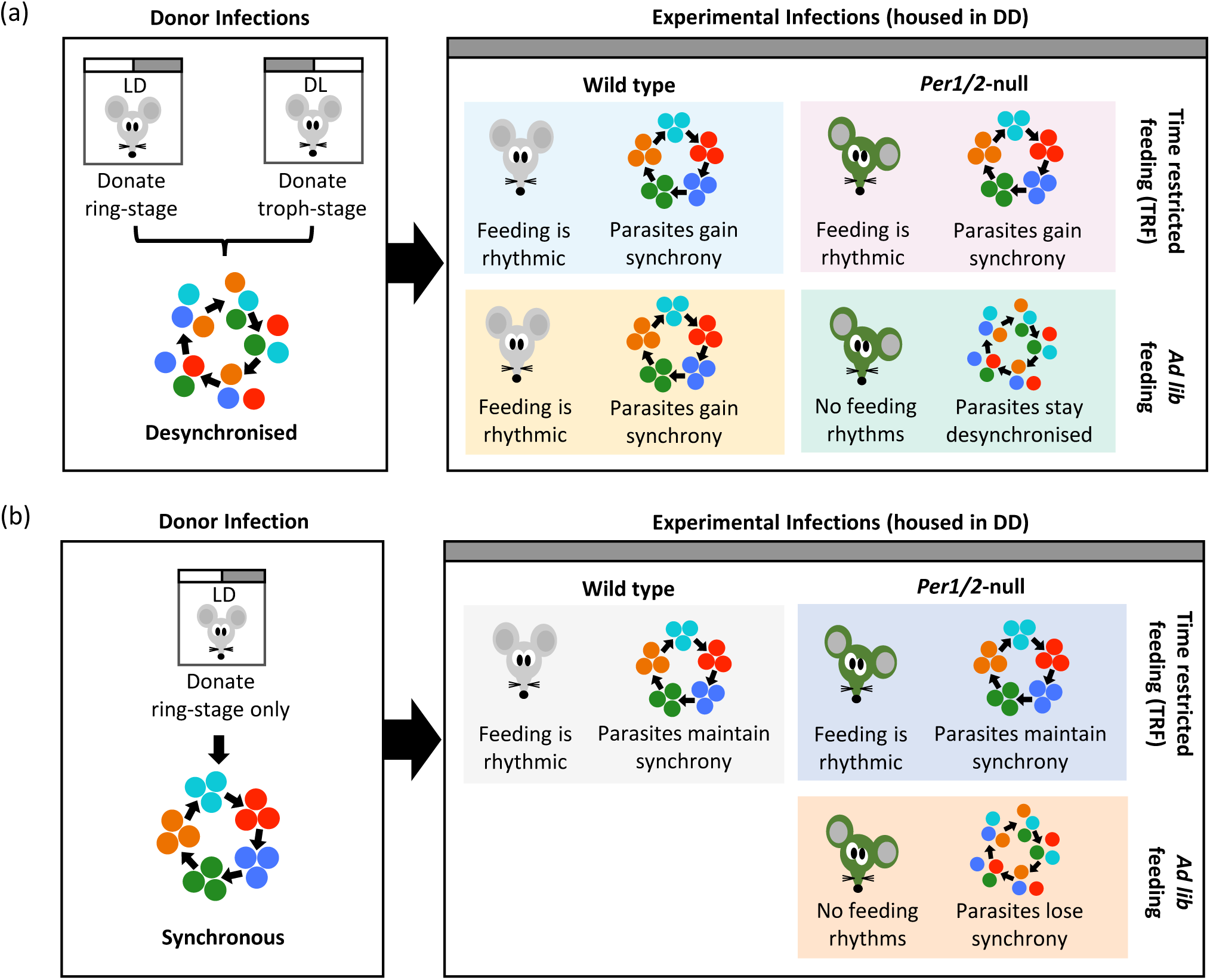
Experimental designs and predictions. Donor infections from mice housed in light:dark and/or dark:light were used to generate a desynchronised (a; ring stage + trophozoite stage parasites) and synchronous (b; ring stage parasites only) inocula for initiating experimental infections. Wild type (WT) or *Per1/2*-null clock-disrupted mice were given constant access to food (*ad lib*) or fed on a Time Restricted Feeding (TRF) schedule in which food access was restricted to only 10h per day. These mice were used as hosts for experimental infections and sampled every 4h for 32h on Day 5 and 6 post infection. We predicted that desynchronised infections will become synchronous in mice in which feeding is rhythmic (both WT groups and *Per1/2*-null TRF) but will remain desynchronised in the *ad lib* fed *Per1/2*-null mice due to a lack feeding rhythms (a). For infections initiated with synchronous parasites, we predicted that parasites will maintain synchrony in WT and *Per1/2*-null TRF groups and that their schedule becomes matched to the timing of host feeding, but that parasites will lose synchrony in *ad lib* fed *Per1/2*-null that lack feeding rhythms (b).

### Parasites and hosts

Hosts were either wild type (WT) C57BL/6J strain or *Per1/2*-null clock-disrupted mice previously backcrossed onto a C57BL/6J background for over 10 generations. *Per1/2*-null mice (kindly donated by Michael Hastings (MRC Laboratory of Molecular Biology, Cambridge, UK), generated by David Weaver (UMass Medical School, Massachusetts, USA)) have an impaired TTFL clock and exhibit no circadian rhythms in physiology and behavior. For example, they exhibit arrhythmic locomotor activity when placed in constant conditions such as constant darkness [33, 34]. All experimental WT and *Per1/2*-null mice were 8-10 weeks old and housed at 21°C in DD (continuous darkness) with constant dim red LED light for the duration of the experiments. Note, donor mice were housed in light-dark cycle conditions to generate synchronous parasites for the initiation of experimental infections. All mice were acclimatized to their feeding treatments (see below) and to DD conditions for 3 weeks before and throughout infections. As the free running period (the time taken to complete one cycle of an endogenous rhythm in the absence of environmental time cues) of our mice is very close to 24hours (23.8-23.9h, electronic supplementary material (ESM)) when placed in DD these mice will exhibit a schedule similar to the initial LD conditions they were raised in. Therefore, we define subjective day (rest phase) for wild type mice as 07:00-19:00 GMT and subjective night (active phase) as 19:00-07:00 GMT. All mice were provided with unrestricted access to drinking water supplemented with 0.05 % para-aminobenzoic acid (to supplement parasite growth). All mice were housed individually to avoid any influence of conspecific cage-mates on their rhythms. On Day 0, each mouse was infected with 5 × 10^6^ *P. chabaudi* (clone DK) parasitized red blood cells administered via intravenous injection. All procedures were carried out in accordance with the UK Animals (Scientific Procedures) Act 1986 (PPL 70/8546).

### Experimental designs

#### Experiment 1 – does a host feeding rhythm restore the IDC schedule of desynchronised parasites?

Wild type WT mice and *Per1/2*-null (n=5 per group) mice were fed on either a time restricted feeding schedule (TRF; fed for 10hours per day) or had access to food *ad libitum* (*ad lib*). This generated 4 treatment groups with the following characteristics (Figure 1a): (i) WT *ad lib* mice with 24h access to food that followed their normal free-running rhythms and primarily fed in their subjective night (19:00-07:00 GMT); (ii) WT TRF mice fed during subjective day (09:00-19:00 GMT) experienced temporal misalignment between rhythms controlled by the suprachiasmatic nucleus (SCN; which free-runs at a timing close to their previous entrainment to LD conditions) and peripheral rhythms, which are driven by feeding [11]; (iii) *Per1/2*-null *ad lib* mice that had continuous access to food and were arrhythmic (ESM) and (iv) *Per1/2*-null TRF mice fed during subjective day that only experienced rhythms resulting from a set daily period for feeding. Infections in all mice were initiated with a population of desynchronised parasites at 08:30 GMT. The inoculum was a 50:50 mix of parasites 12 hours apart in the IDC. Specifically, ring stages (donated from donors in a 12:12 light:dark cycle (LD)) and late trophozoite stages (donated from dark:light (DL) donors)(Figure 1a).

#### Experiment 2 – do parasites lose IDC synchrony in the absence of a host feeding rhythm?

We generated 3 groups of n=5 mice (Figure 1b): (i) WT TRF mice fed during subjective day; (ii) *Per1/2*-null TRF mice fed during subjective day; and (iii) *Per1/2*-null *ad lib* food. Infections in all mice were initiated with a population of synchronous ring stage parasites collected at 08:30 GMT from a single donor in a 12:12 light:dark cycle (LD). This meant the parasites entered experimental hosts early in the feeding period, which is 12 hours out of phase to when rings stages peak in control infections (Figure 1b). Generating a mismatch between incoming parasites and the hosts feeding rhythm tests whether the IDC becomes rescheduled to match the feeding rhythm in the TRF groups, avoiding an outcome of the ICD being unable to change obscuring the interpretation of results, following Prior et al 2018 [11].

#### Sampling & data collection

All experimental mice were sampled at 4-hourly intervals for 32 hours beginning at 08:00 (GMT) Day 5 to 16:00 Day 6 post infection (pi). Previous work [11] revealed that synchronous parasites in infections mismatched to the host’s feeding rhythm by 12 hours exhibit a rescheduled IDC within four days, and this is verified by the rescheduling of the IDC in the WT TRF mice fed during subjective day in experiment 2 (Figure 2a; Figure 3a). At each sampling point, blood was collected from the tail vein and parasites at each IDC stage quantified from thin blood smears. Stages were characterised by morphology, based on parasite size, the size and number of nuclei and the appearance of haemozoin (as per [11] and [2]). Red blood cell (RBC) densities were measured at each sampling time by flow cytometry (Z2 Coulter Counter, Beckman Coulter). Mouse weights were measured on Day 2 PI and Day 6 PI at 16:00 GMT. All procedures were carried out in dim red LED light. Before infection, we verified that locomotor activity (movement around the cage) and internal body temperature of WT mice followed the free-running rhythms expected in DD given their previous entrainment to LD, and whether locomotor activity can be used as a good proxy for feeding events (ESM).

**Figure 2.**
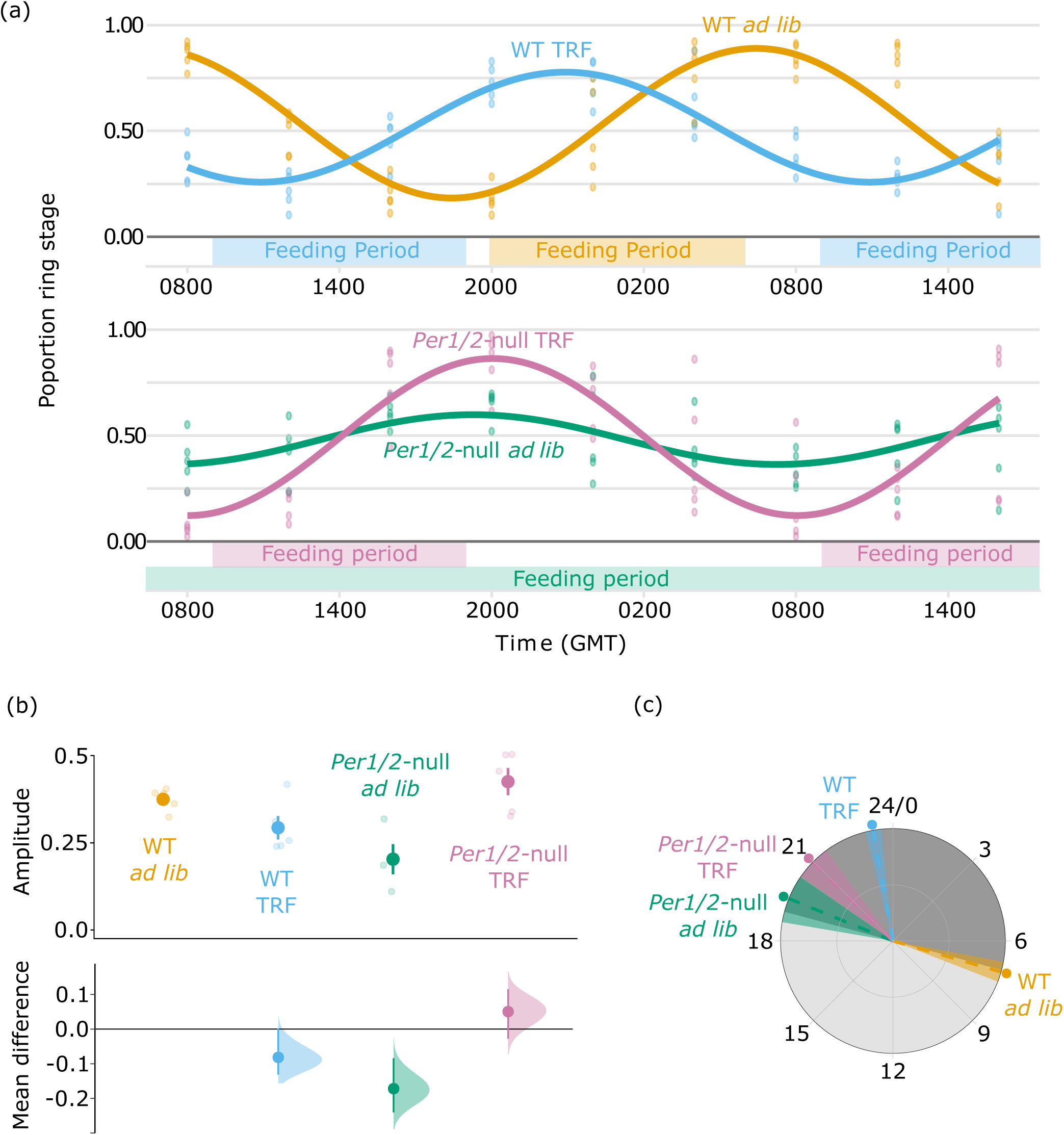
The IDC of initially desynchronised parasites becomes coordinated to host feeding rhythms. Population cosinor model fits and data points from each individual infection (a). Amplitude (b) and phase in hours (GMT) (c) were calculated from cosinor model fits from each individual mouse (lighter points) and then summarised as a mean ±SEM, points with error bars in (b), and circular mean ±SD point with dashed line and shading in (c). For amplitude (b), effect sizes relative to the ‘WT *ad lib*’ group are plotted on the lower axes as a bootstrap sampling distribution (mean difference ± 95% CI depicted as a point with error bars). For all parts, infections in Wild Type (WT) hosts are coloured orange and blue, and infections in *Per1/2*-null mice are coloured green and purple. TRF indicates ‘Time Restricted Feeding’ with food only available for 10 hours each day (feeding period indicated above x axis in (a)). Grey shading in (c) represents active (dark shading; 19:00-07:00) and rest (light shading; 07:00-19:00) periods relative to wild type mice in DD.

**Figure 3.**
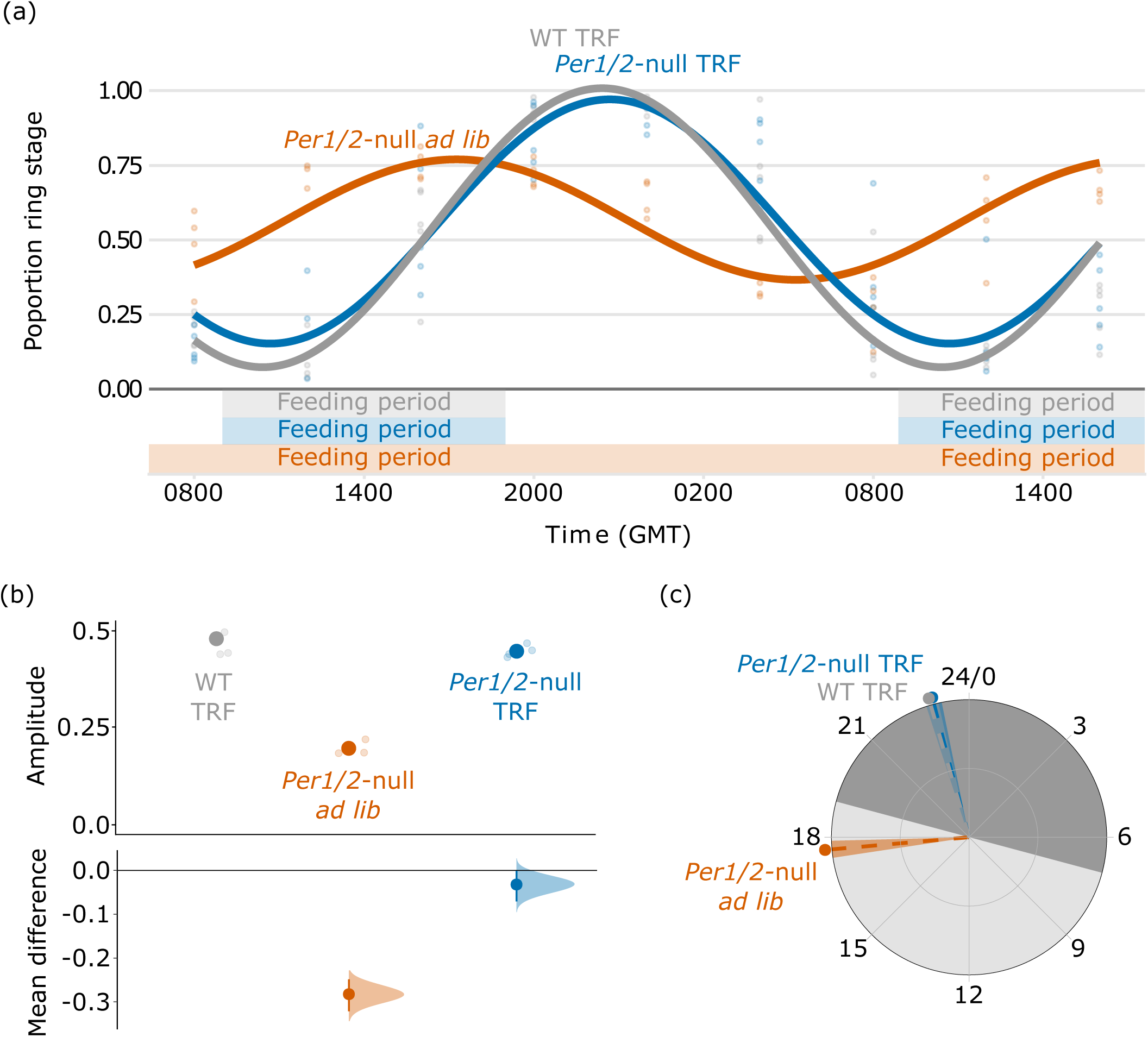
The IDC loses synchrony in hosts without feeding rhythms. Population cosinor model fits and data points from each individual infection (a). Amplitude (b) and phase in hours (GMT) were calculated from cosinor model fits from each individual mouse (lighter points) and then summarised as a mean ±SEM, points with error bars in (b), and circular mean ±SD point with dashed line and shading in (c). For amplitude (b), effect sizes relative to ‘WT TRF’ group are plotted on the lower axes as a bootstrap sampling distribution (mean difference ± 95% CI depicted as a point with error bars). For all parts, infections in Wild Type (WT) are coloured grey, and infections in *Per1/2*-null mice are coloured orange and blue. TRF indicates ‘Time Restricted Feeding’ with food only available for 10 hours each day (feeding period indicated above x axis in (a)). Grey shading in (c) represents active (dark shading; 19:00-07:00) and sleep (light shading; 07:00-19:00) periods relative to wild type mice in DD.

#### Statistical Analysis

Time of onset of locomotor activity and overall period estimates for locomotor activity and body temperature data were calculated using Clocklab (Actimetrics). Harmonic regression (periodicity analysis) was performed on parasite data using Circwave (v. 1.4, courtesy of R. Hut; http://www.euclock.org). All other statistical analyses were carried out using R version 3.5.0. Correlations between feeding events and locomotor activity was tested using a generalised linear model with a zero-inflated poisson regression (package pscl). Output model fit measures for amplitude and phase, as well disease severity (red blood cell loss) and parasite performance (maximum parasite density) measures, were compared between groups using general linear models and models were confirmed for best fit by comparing AICC values. All models met model assumptions: independence of data points, normality of residuals and homogeneity of variances (confirmed through assessing the model plots, the Shapiro–Wilk test and Bartlett’s test). Whilst some variation in parasite density was observed across treatment groups, any underlying differences in replication rate are not large enough to introduce biases associated with estimating synchrony from IDC stage proportion data [35]. Mean effect sizes were calculated using bootstrapped coupled estimation (package dabestr). Measures of red blood cell loss were calculated by taking the maximum RBC value across the time series and subtracting the minimum RBC value. This minimised any issues associated with comparing data for the same GMT across groups in which host and parasite rhythms may be phased differently. For measures of circadian phase, Bayesian circular GLM’s (selected using DIC) were used (package circglmbayes), which allows regressing a circular outcome on linear and categorical predictors.

## RESULTS

### Assumptions of the experimental designs

We first verified that only WT mice exhibit rhythms in locomotor activity and body temperature and confirmed arrhythmic activity of *Per1/2*-null *ad lib* mice (ESM). Second, we reveal that locomotor activity can be used as a proxy for feeding rhythms in *Per1/2*-null TRF mice (ESM).

### Experiment 1 – does a host feeding rhythm restore the IDC schedule of desynchronised parasites?

We compared IDC rhythms in terms of synchronicity (amplitude) and timing (phase) of the proportion of parasites at ring stage (a morphologically distinct ‘marker’ stage after which all other parasite stages follow in a predictable manner)[11]. By day 5-6 post infection, the IDC of parasites in all WT mice and *Per1/2*-null TRF mice had become scheduled to coincide with host feeding rhythms (Figure 2a). Amplitude differed significantly between groups (Figure 2b; genotype:feeding_regime: *χ*^2^_15_= 0.43, *p* < 0.001). Specifically, parasites in *Per1/2*-null TRF mice had the highest amplitudes (mean ±SEM: 0.85 ± 0.08) followed by wild type *ad lib* infections (0.75 ± 0.03), and then wild type TRF infections (0.59 ± 0.07), with *Per1/2*-null *ad lib* infections (0.41 ± 0.09) exhibiting approximately half the amplitude of parasites in hosts with feeding rhythms. We also found differences in the timing of peak proportion ring stages between groups explained by a host genotype:feeding_regime interaction (Figure 2c; ESM Table S1). Rings in WT mice peaked at the end of their hosts subjective feeding regime (circular mean ±SD (hours GMT): WT *ad lib* = 7.06 ± 0.35) while parasites in TRF mice peaked within 1-2 hours of end of their restricted feeding regime (WT TRF = 23.32 ± 0.27, *Per1/2*-null TRF = 20.96 ± 0.61). Parasites in *ad lib* fed *Per1/2*-null mice peaked at 19.47 (± 0.83). See ESM Table S2 for a summary of bootstrapped mean effect sizes.

We also assessed whether anaemia and parasite performance varies according between WT and *Per1/2*- null mice. Neither host genotype, feeding regime or their interaction significantly affected RBC loss (genotype:feeding_regime: *χ*^2^_16_= 1.82 × 10^17^, *p* = 0.60; host genotype: *χ*^2^_17_= 1.06 × 10^17^, *p* = 0.69; feeding regime: *χ*^2^_18_= 1.27 × 10^18^, *p* = 0.15), with hosts losing an average of 2.50 ±0.18 SEM × 10^9^ ml^−1^ RBCs during the sampling period (ESM Figure 3a; ESM Table S2). In contrast, host genotype had a significant effect on maximum parasite density (*χ*^2^_17_= 1.15 × 10^18^, *p* = 0.002) in which parasites infecting WT hosts achieved maximum densities ∼40% higher than those parasites infecting *Per1/2*-null mice (mean ±SEM × 10^9^ml^−1^: WT = 1.69 ±0.08, *Per1/2*-null = 1.22 ±0.10; ESM Figure 3b; ESM Table S2). Neither feeding regime (*χ*^2^_17_= 2.16 × 10^16^, *p* = 0.63) nor it’s interaction with host genotype (*χ*^2^_16_= 3.42 × 10^16^, *p* = 0.55) had an effect on maximum parasite density.

### Experiment 2 – do parasites lose IDC synchrony in the absence of a host feeding rhythm?

By day 5-6 post infection, the IDC of parasites in all TRF fed mice had rescheduled to coincide with host feeding rhythms (Figure 3a). Amplitude differed significantly between host feeding regimes (Figure 3b; *χ*^2^_9_= 0.66, *p* < 0.001). Parasites in TRF mice exhibited high amplitudes (mean ±SEM: WT TRF = 0.96 ± 0.04, *Per1/2*-null TRF = 0.90 ± 0.02) whereas amplitudes for parasites in *Per1/2*-null *ad lib* mice were more than 50% lower (0.39 ± 0.02). These differences were not influenced by host genotype (*χ*^2^_9_= 0.01, *p* = 0.11). In addition, we found differences in IDC timing that are explained by host feeding regime (Figure 3c; ESM Table S1). Parasites in TRF mice peaked 4 hours after their host’s feeding period (circular mean ±SD (hours GMT); WT TRF = 22.92 ± 0.22, *Per1/2*-null TRF = 23.01 ± 0.22) whereas parasites in *Per1/2*-null *ad lib* mice peaked 8 hours earlier than TRF groups (17.66 ± 0.24). Again, host genotype did not explain these differences (ESM Table S1). See ESM Table S2 for a summary of bootstrapped mean effect sizes.

Red blood cell loss was influenced by both host genotype (*χ*^2^_11_= 4.17 × 10^18^, *p* < 0.001) and host feeding regime (*χ*^2^_11_= 4.56 × 10^17^, *p* = 0.025; ESM Figure 4a; ESM Table S2). Specifically, WT TRF lost the most RBC (30-57% more than both groups of *Per1/2*-null mice), and *ad lib* fed *Per1/2*-null mice lost 20% more RBC’s than their *Per1/2*-null TRF counterparts (mean ±SEM × 10^9^ml^−1^: WT TRF = 3.57 ± 0.13, *Per1/2*-null TRF = 2.28 ± 0.15, *Per1/2*-null ad lib = 2.73 ± 0.13). Host genotype alone also had an effect on maximum parasite density (genotype: *χ*^2^_2_= 4.51 × 10^17^, *p* = 0.002; feeding regime: *χ*^2^_11_= 9.36 × 10^14^, *p* = 0.89) with parasites in WT TRF hosts achieving maximum parasites densities 24% higher than parasites in *Per1/2*- null mice (mean ±SEM × 10^9^ml^−1^: WT TRF = 1.89 ± 0.11, *Per1/2*-null = 1.51 ± 0.07; ESM Figure 4b; ESM Table S2).

## DISCUSSION

Our results demonstrate that timing and synchrony of the IDC of the malaria parasite *P. chabaudi* is not dependent on rhythms driven by the canonical “TTFL” circadian clock of hosts, and that feeding rhythms alone establish the IDC schedule. The first experiment revealed that infections in wild type hosts initiated with desynchronised parasites became synchronized and that the timing of IDC transitions became coordinated with the timing of feeding (ring stages peaking 2-4 hours after food intake ends), consistent with Prior *et al* [11]. Strong IDC rhythms also emerged in hosts that do not possess a functioning TTFL clock but do have enforced feeding rhythm with very similar timing (reaching an average peak ring stage proportion of 86% within an hour after the feeding period ends). Furthermore, in *ad lib* fed clock-disrupted hosts, which feed in many small irregular bouts across each 24hrs, the IDC remained desynchronised (ring stage proportion remaining at around 50% across all sampling points). Consistent with these phenomena, our second experiment revealed that infections initiated with synchronous parasites remained synchronous and became coordinated to the timing of host feeding in both wild type and clock disrupted mice with feeding rhythms. Whereas in *ad lib* fed clock-disrupted mice, the IDC rhythm became dampened (peak in ring stages dropping from 100% to 75%). Put another way, we show that an IDC schedule emerges in hosts with a feeding rhythm independently of whether they have a TTFL clock, and the IDC schedule is lost in hosts without a feeding rhythm. Whilst the IDC rhythm of synchronous infections of *ad lib* fed clock-disrupted mice became dampened, it did not become fully desynchronised. There are two non-mutually exclusive reasons for this. First, there are likely to be development constraints acting on the duration of each IDC stage and the overall IDC length. If the minimum and maximum duration of the IDC is close to 24 hours, or stage durations similarly constrained, natural variation in duration will take more cycles to erode synchrony than in our experiment. Second, even in completely asynchronous infections, the expansion of parasite number due to each asexual stage replacing itself with multiple progeny, can generate the illusion of strong synchrony [35].

Why should host feeding rhythms set the schedule for the IDC? Blood glucose concentration follows a daily rhythm and has been implicated as the specific factor responsible for the timing of the IDC schedule. Glucose tolerance oscillates across the day in a circadian manner and behavioural factors, such as activity, feeding and fasting, strongly affect glucose metabolism. This makes it difficult to quantify the independent effect of the circadian system on the 24h diurnal variations in glucose concentration [36]. Glucose regulation is a tightly controlled process, achieved via the antagonistic effects of the hormones insulin and glucagon, and involves the contribution of several different organs (liver, pancreas) to dampen perturbations due to feeding and fasting. Malaria infection disrupts this balance by exacerbating the drop in blood glucose concentration at the end of the active phase [10]. Perhaps this drop, or the rapid rise in blood glucose concentration at the start of the feeding period, could either signal time-of-day to parasites and/or enforce a schedule in which only the most glucose demanding IDC stages (late trophozoites and schizonts) can develop during the feeding period. However, other resources required by later IDC stages, including amino acids essential for protein synthesis [37], purines (in particular hypoxanthine) for nucleic acid synthesis, and lysophosphatidylcholine (lysoPC) for various processes such as cell membrane production [38], are also likely derived by the parasite from the host’s food. Feeding-driven rhythms in these factors (in addition to/instead of blood glucose concentration) may ultimately drive the IDC schedule. Because glucose – and other metabolites derived from food – are generally elevated throughout the feeding period, which is a significant proportion of the circadian cycle, late IDC stage parasites have a long window in which they can develop. Given that the timing of the IDC schedule appears more precise than simply IDC completion occurring at some point during the feeding period, we suspect malaria parasites are, at least in part, capable of organizing the IDC schedule themselves to orient it to the timing of incoming resources.

Whilst our results strongly implicate food rhythms as responsible for the IDC schedule, other rhythms remain as possibilities. First, circadian rhythms independent of the TTFL could operate in clock-disrupted mice and schedule the IDC. For example, lipid levels in hepatic cells maintain oscillations in clock-disrupted mice (specifically *Bmal1* knockouts) albeit in a different phase to wild type mice [39], and under specific experimental conditions, food anticipatory activity can be observed in other clock-disrupted mice [40]. The mechanisms of these rhythms are not fully understood but daily oxidation-reduction rhythms exist (which occur within mammalian RBCs [41] and are evolutionarily conserved [42]) and may be linked to cellular flux in magnesium ions [43]. Whether such rhythms are reliable time-cues during infection or if they can be entrained by the timing of feeding are unknown. Second, in small mammals, body temperature rhythms are influenced by a combination of the circadian clock, metabolism, and locomotor activity. Temperature rhythms can entrain host cells (including RBCs) and other parasites (e.g. *Trypanosoma brucei*) [9], and malaria parasites do respond to temperature change (e.g. when taken up by ectothermic mosquitoes) [44, 45]. However, it is unlikely that the IDC schedule is set by a temperature rhythm. Prior *et al* (2018) reveal inverted IDC rhythms in day- and night-fed mice but host temperature rhythms are not inverted. Similarly, both the *ad lib* fed and TRF fed wild type hosts in the experiments presented here exhibit daily temperature rhythms that only differ in onset by a few hours, but the IDC schedule differs by 8 hours. Third, a combination of host rhythms may impact on the IDC schedule, including minor contributions from non-feeding rhythms. This may explain why the degree of synchronicity is slightly reduced in TRF wild type hosts compared to *ad lib* fed wild type hosts, and highest in TRF clock-disrupted mice: the separation of food-entrained and SCN-driven rhythms in TRF wild type hosts may result in conflicting time-cues, whereas the lack of TTFLs in all cells in the clock-disrupted mice may mean that only food-related time-cues are present.

We were also able to use our data to test whether parasite performance is enhanced in clock-disrupted hosts, due perhaps to lack of regulation/coordination of clock-controlled immune responses [29, 32]. However, we find that the maximum parasite density is approximately 25 – 40% (across both experiments) lower in infections of clock disrupted compared to wild type hosts. Clock disruption might reduce the ability of hosts to process and metabolise food efficiently, making these hosts a poorer resource for parasites. For example, PER1 and PER2 have a regulatory role in the circadian control of heme synthesis, a primary resource for parasites [46] and loss of PER2 is implicated in making RBC more susceptible to oxidative stress, decreasing levels of ATP and shortening RBC lifespan [47]. Further, clock disruption affects host nutrition via an interplay with microbiota [48]. Parasite performance is linked to host nutrition because caloric restriction leads to reduced parasite densities [49]. However, if either clock disruption and/or our time-restricted-feeding regime caused caloric restriction we would expect this to manifest as clock disrupted mice – especially in the TRF groups – as having the lowest weights or the greatest weight loss. In contrast, *Per1/2*-null TRF mice were the heaviest in experiment 1 and *Per1/2*-null *ad lib* mice lost the most weight in experiment 2 (ESM Table S3). Another, non-mutually exclusive, possibility is that the IDC becomes scheduled to coordinate with feeding rhythms faster in wild type mice due to facilitation by a host clock-controlled process and the faster parasites can reschedule, the lower the costs of being uncoordinated to the host rhythm. However, that parasite performance does not differ between infections remaining / becoming desynchronised versus synchronous within the same type of host (i.e. *Per1/2*-null) suggests either that there are no major costs to parasites of being desynchronised or that it is advantageous for them to match the degree of rhythmicity of their host. For example, a desynchronised IDC in arrhythmic hosts (*Per1/2*-null *ad lib* fed) may be the best strategy to avoid exploitation competition between parasites for arrhythmic resources. Whilst the costs of virulence, as measured by weight loss, do not appear to differ between wild type and clock-disrupted hosts, the findings for RBC loss are more complicated (and do not clearly mirror maximum parasite densities). No significant difference between feeding regimes or host genotypes were detected when infections were initiated with desynchronised parasites. But, in infections initiated with synchronous parasites, wild type hosts became the most anaemic and clock-disrupted hosts with a feeding rhythm lost the least RBC. Further work is needed to establish whether a loss of canonical clock regulation affects the ability of host to control or tolerate parasites.

## CONCLUSION

The schedule (timing and synchrony) of the malaria parasite’s IDC is not reliant on a functioning host canonical circadian clock. The speed with which the IDC schedule changes, its precision, and the modest loss of parasite number involved in rescheduling [13], suggest the parasite is actively aligning certain developmental stages with host feeding rhythms to take advantage of periodicity in a resource(s). The precise cue(s) that parasites use to schedule the IDC is unknown but we propose it is directly related to feeding events, and not associated with the food-entrained peripheral TTFL in the organs or the central oscillator in the SCN. Our data also highlight a complex interplay between host rhythms, features of the IDC schedule, parasite fitness (as approximated by maximum density), and disease severity. Unravelling these complexities may reveal measures to minimize disease severity and improve recovery, whilst reducing parasite fitness.

## Supporting information

ESM Figure S1

ESM Figure S2

ESM

ESM Figure S3

ESM Figure S4

## ETHICS

All procedures were carried out in accordance with the UK Animals (Scientific Procedures) Act 1986 (PPL 70/8546).

## DATA ACCESSIBILITY

The datasets supporting the conclusions of this article are available in the Edinburgh DataShare repository, https://doi.org/10.7488/ds/2622

## AUTHOR’S CONTRIBUTIONS

AOD and SER conceived the study, AOD and KP carried out the experiments, AOD analysed the data, and all authors wrote the manuscript.

## ACKNOWLEDGEMENTS

We thank Ronnie Mooney for technical assistance and Giles K.P. Bara for advice.

