## Supplementary figures and images for "Host circadian clocks do not set the schedule for the within-host replication of malaria parasites"

### ESM Figure S1

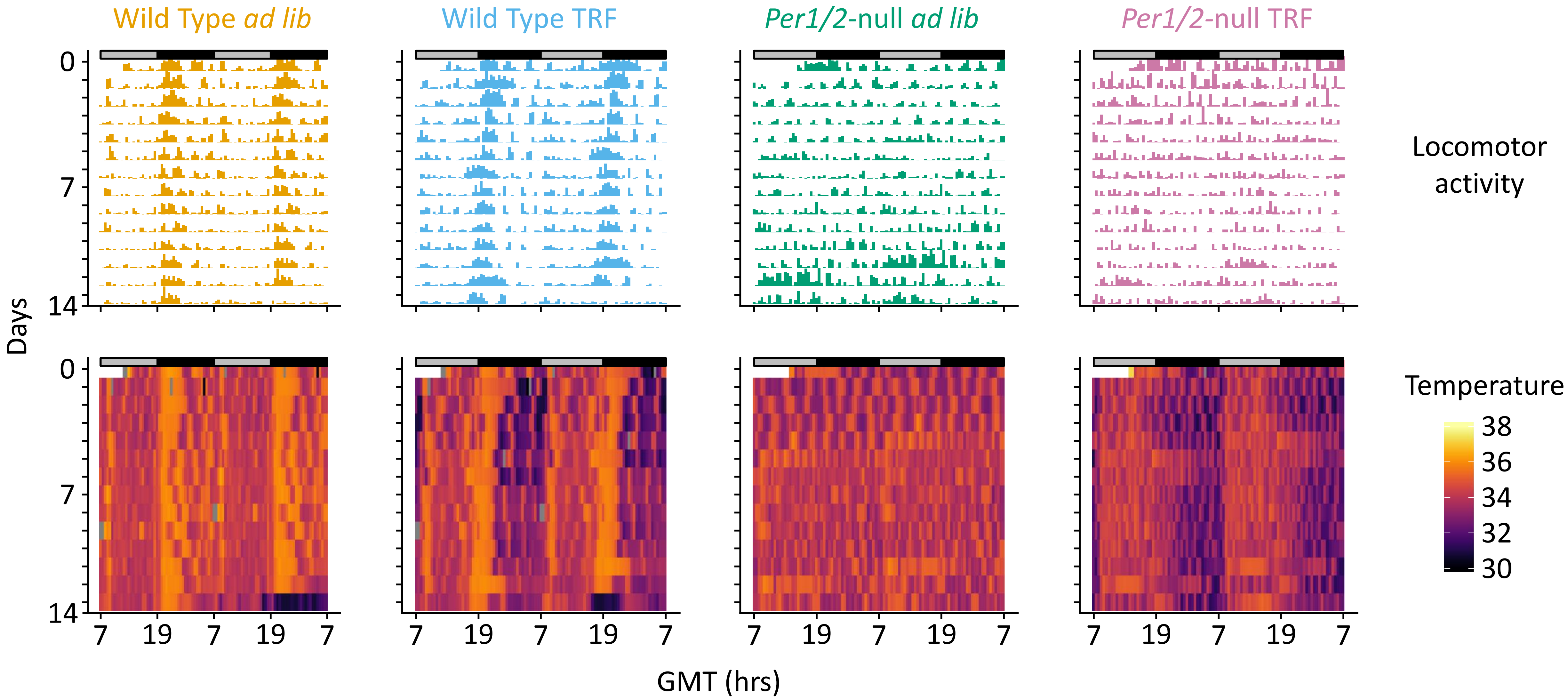

### ESM Figure S2

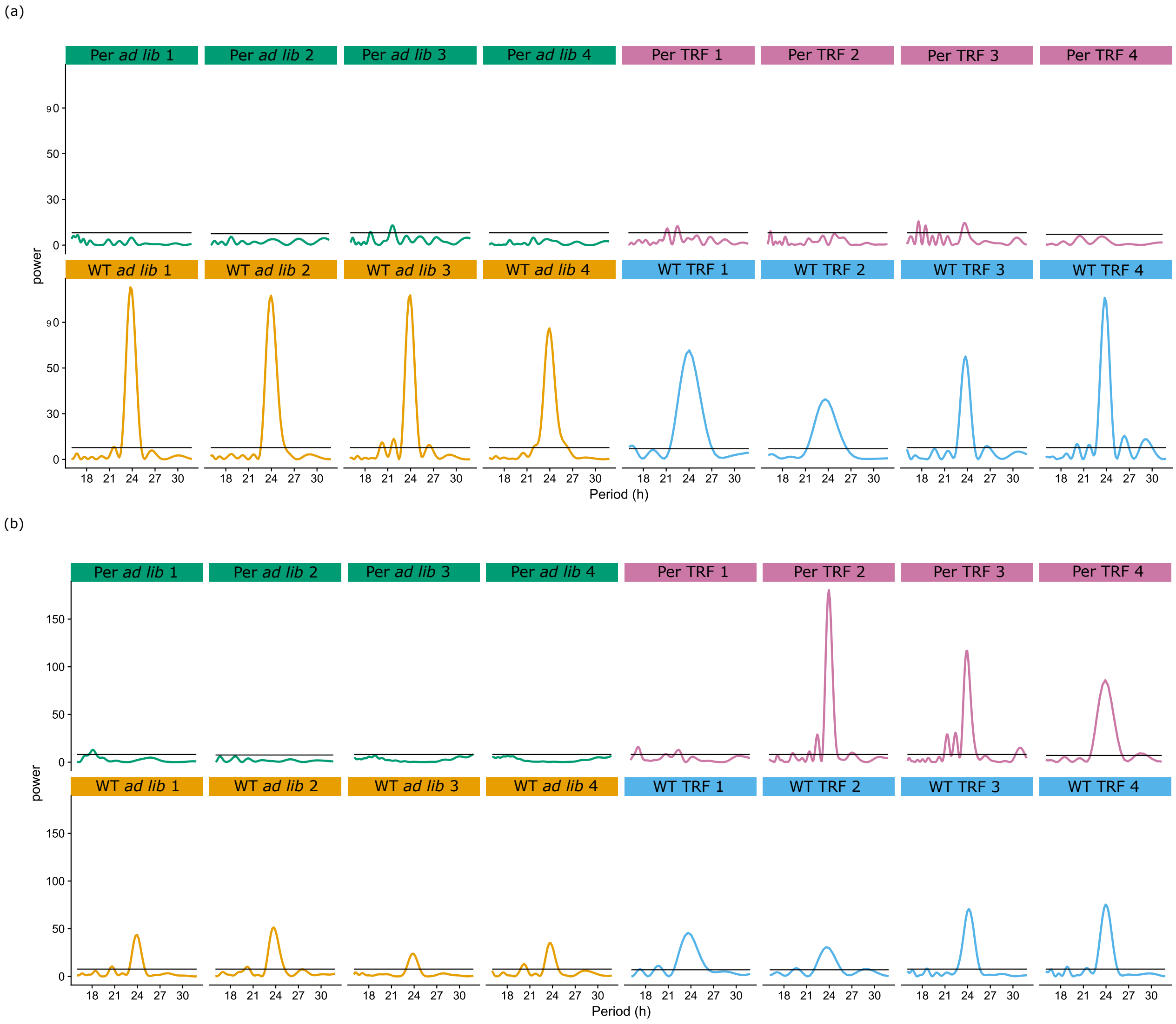

### ESM Figure S3

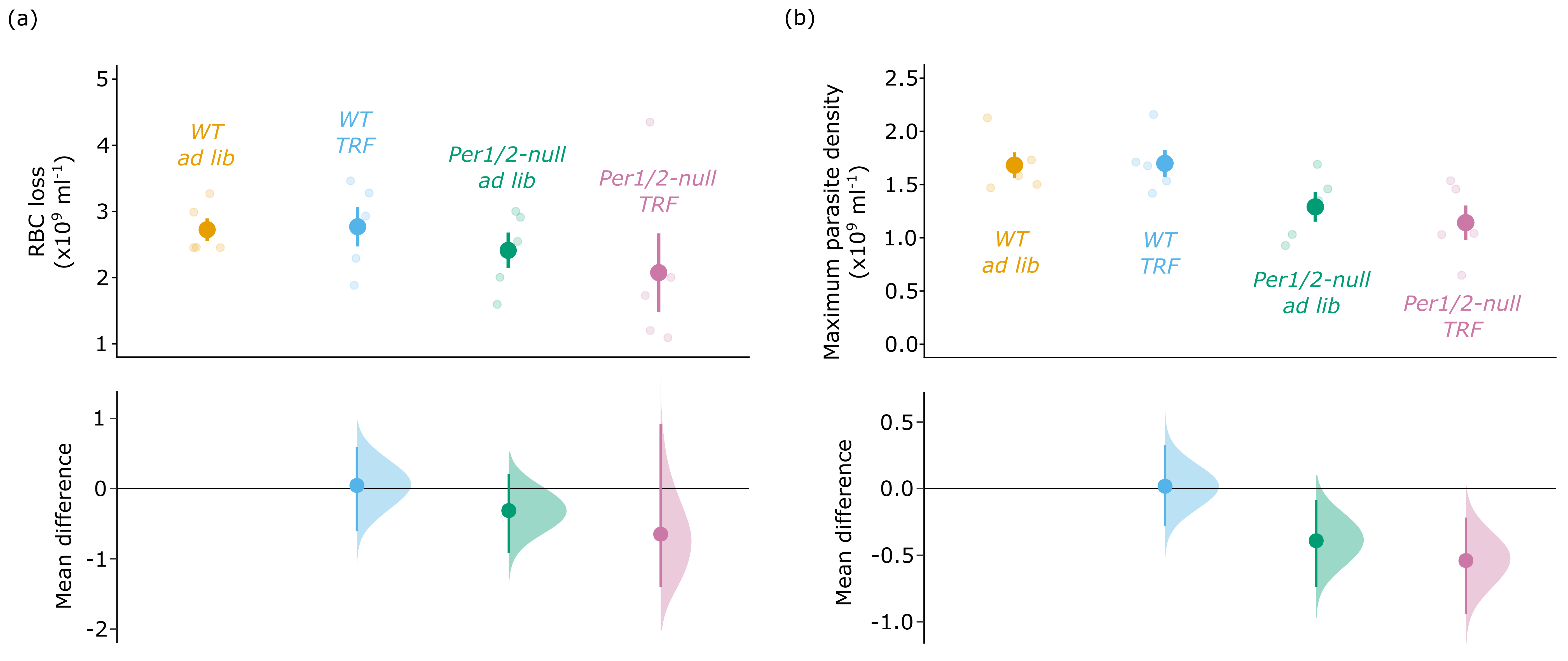

### ESM Figure S4

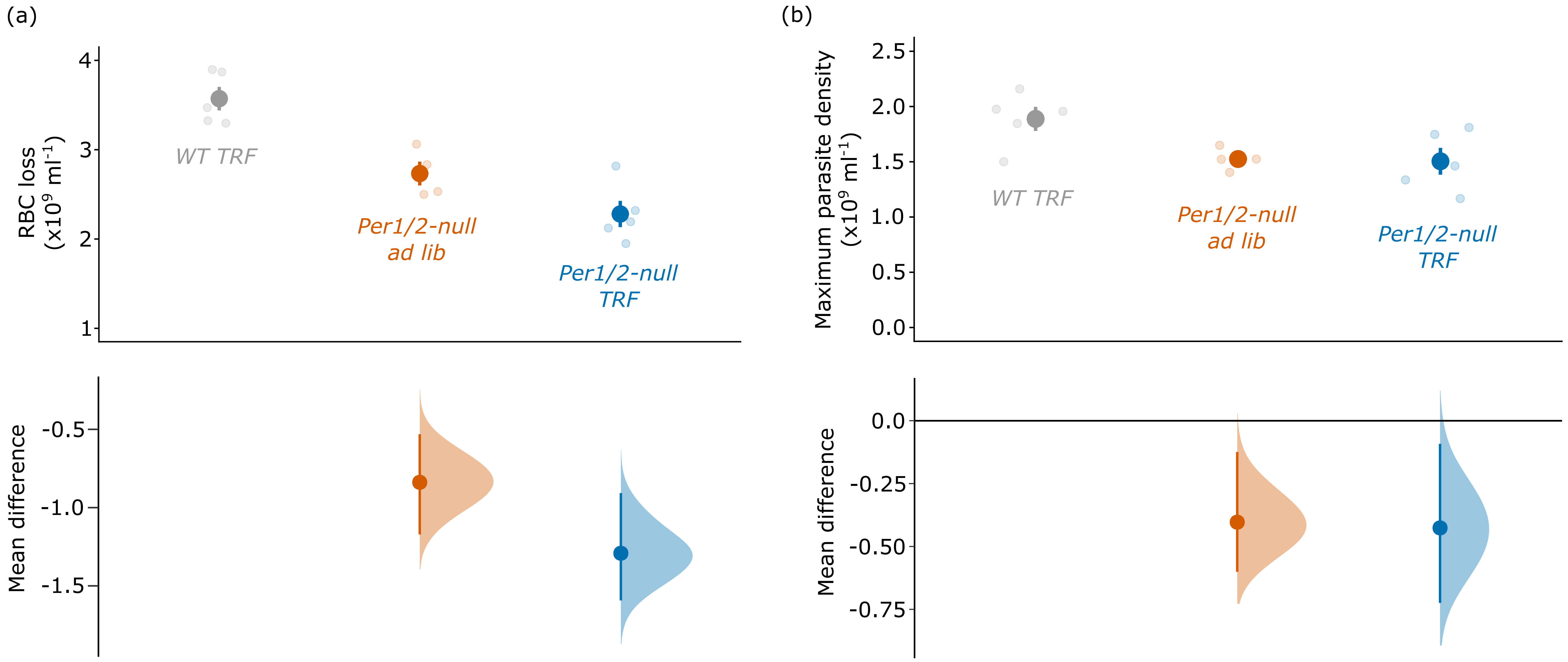
