## Supplementary material for "Host circadian clocks do not set the schedule for the within-host replication of malaria parasites": ESM

**ELECTRONIC SUPPLEMENTARY MATERIAL**

**METHODOLOGY**

***Locomotor and internal body temperature data collection***

We measured activity and temperature pre- and during infection without disturbance to the animals using the Actual Home Cage Analysis system (Actual HCA, Actual Analytics Ltd, Edinburgh, Scotland) [1]. Specifically, Biotherm13 RFID tags (radio frequency identification; Biomark, Idaho, USA) were injected subcutaneously and both positional and temperature readings were recorded every 10ms (and then summed into 1 minute bins) via an array of antennas (spaced ~11cm apart) under the cage.

To determine if locomotor activity can be used as a good proxy for feeding events, two groups of five uninfected WT and *Per1/2*-null mice were group housed (separated by genotype) in DD and filmed for 24 hours. Feeding bouts were classified from observing mice nibbling on food pellets from the hopper. These data were summed to 15 minute bins and compared with locomotor activity data obtained from RFID tags and the Actual Home Cage Analysis system.

**RESULTS**

***Assumptions of the experimental designs***

*Rhythms in locomotor activity*

Only wild type mice exhibited daily periodicity in locomotor activity with a mean period of 23.96 (± 0.01 SEM) for *ad lib* fed and 23.82 (± 0.03 SEM) for TRF fed (Figure S1; Figure S2). WT *ad lib* fed mice had an average median onset of activity per day at 20.1 (±0.21 SEM; GMT) while TRF WT mice started their activity 3 hours earlier at 17.2 (±0.54 SEM; GMT; χ^2^_1_= 4.03, *p* < 0.001). Level of activity during rest periods compared to active periods in WT mice was not influenced by feeding regime (χ^2^_381_= 21.78, *p* = 0.72). *Per1/2*-null mice did not exhibit rhythmic locomotor activity (i.e. are behaviourally arrhythmic) as evident from actograms (Figure S1) and lomb-scargle periodograms (Figure S2). Total activity per 24h period was 41% lower in *Per1/2*-null mice compared to WT mice (mean transitions ±SEM: WT = 1636 ±70.7, *Per1/2*-null = 958 ±48.5; χ^2^_270_= 30.28 x 10^6^, *p* < 0.001) and activity was not influenced by feeding regime (χ^2^_269_ = 0.56 x 10^6^, *p* = 0.26) or its interaction with genotype (χ^2^_268_= 0.01 x 10^6^, *p* = 0.86).

*Rhythms in internal body temperature*

Both WT TRF and *ad lib* mice exhibited rhythms in internal body temperature (becoming ~1°C cooler during rest periods; Figure S1) with free-running periods close to 24 hours (mean period ± SEM: WT *ad lib* = 23.96 ± 0.01, WT TRF = 21.74 ± 2.1). Unlike locomotor activity, *Per1/2*-null TRF mice exhibited rhythmicity in body temperature with a mean period of 26.26 (±5.97 SEM), becoming, ~1°C cooler during periods when food was unavailable. No such temperature rhythms were observed in *ad lib* fed *Per1/2*-null mice (Figure S1; Figure S2).

*Locomotor activity as a proxy for feeding activity*

Finally, the experimental designs required that locomotor activity was strongly correlated with feeding events in both WT mice and *Per1/2*-null mice, which was the case (Zero-inflated count model; WT: z = 21.22, *p* < 0.001, *Per1/2*-null: z = 16.71, *p* < 0.001). Non-feeding events were also correlated with zero activity (Zero-inflated binomial model; WT: z = -2.54, *p* = 0.011, *Per1/2*-null: ZeroInfl binomial model: z = -2.770, *p* = 0.006). If animals were moving they generally were also eating, and therefore locomotor activity can also be used as a proxy for feeding events during periods when food was available.

**SUPPLEMENTARY FIGURES**

**Figure S1**

**Locomotor activity and internal body temperature.** Locomotor activity (top) and internal body temperature (bottom) measures from ‘representative mice’ recorded via subcutaneous RFID chips and an antennae array over 14 days pre-infection (summed into 5 minute bins and double plotted). Light and grey horizontal bars represent subjective day and night, respectively. WT and *Per1/2*-null mice were given food either *ad lib* or had access to food restricted to 10 hours each day between 09:00 and 19:00 (time restricted diet, TRF).

**Figure S2**

**Lomb-scargle periodograms for (a) locomotor activity and (b) body temperature.** Locomotor activity (a) and internal body temperature (b) measures from four ‘representative mice’ per treatment recorded via subcutaneous RFID chips and an antennae array over 14 days pre-infection (summed into 5 minute bins). Dark horizontal line indicates a significance threshold of 0.01. Wild type (WT) and *Per1/2*-null (Per) mice were given food either *ad lib* diet or had access to food restricted to 10 hours per day (time restricted diet, TRF).

**Figure S3**

**Virulence and performance of infections initiated with desynchronised parasites.** (a) Virulence as measured by mean (±SEM) red blood cell (RBC) loss across infections in wild type and *Per1/2*-null mice, according to feeding regime. (b) Parasite performance as measured by mean (±SEM) maximum parasite density. For each, individual infections are shown as lighter points. Effect sizes relative to ‘WT *ad lib*’ group are plotted on the lower axes as a bootstrap sampling distribution (mean difference ± 95% CI depicted as a point with error bars). WT and *Per1/2*-null mice were given food either *ad lib* diet or had access to food restricted to 10 hours per day (time restricted diet, TRF).

**Figure S4**

**Virulence and performance of infections initiated with synchronous parasites.** (a) Virulence as measured by mean (±SEM) red blood cell (RBC) loss across infections in wild type and *Per1/2*-null mice, according to feeding regime. (b) Parasite performance as measured by mean (±SEM) maximum parasite density. For each, individual infections are shown as lighter points. Effect sizes relative to ‘WT TRF’ group are plotted on the lower axes as a bootstrap sampling distribution (mean difference ± 95% CI depicted as a point with error bars). WT and *Per1/2*-null mice were given food either *ad lib* diet or had access to food restricted to 10 hours per day (time restricted diet, TRF).

|  | model | DIC | ΔDIC |
| --- | --- | --- | --- |
| Expt. 1 | **phase ~ genotype:feeding_regime** | **40.27** | **27.84** |
|  | phase ~ genotype + feeding_regime | 55.37 | 12.74 |
|  | phase ~ genotype | 59.93 | 8.18 |
|  | phase ~ feeding_regime | 64.32 | 3.79 |
|  | phase ~ 1 | 68.11 | - |
| Expt. 2 | **phase ~ feeding_regime** | **4.07** | **23.78** |
|  | phase ~ genotype + treatment | 6.32 | 21.53 |
|  | phase ~ genotype | 27.49 | 0.36 |
|  | phase ~ 1 | 27.85 | - |

**Table S1**

**Differences in DIC (ΔDIC) between Bayesian circular generalized linear models explaining parasite phase.** Minimum model in bold. Parasite phase is best explained by a mouse genotype:feeding_regime interaction in Experiment 1 and by feeding regime alone in Experiment 2.

|  | Group | Mean Amplitude difference  (95% CI) | Mean Phase difference (hours) (95% CI) | Mean RBC loss difference (x10^8^ ml^-1^) (95% CI) | Mean max parasite density difference (x10^8^ ml^-1^) (95% CI) |
| --- | --- | --- | --- | --- | --- |
| **Expt. 1** | WT *ad lib*  (reference group) | Reference group | Reference group | Reference group | Reference group |
|  | WT TRF | -0.16  (-0.26, 0.00) | **-7.74  (-9.26, -6.30)** | 0.05  (-0.61 0.60) | 0.17  (-2.81, 3.25) |
|  | *Per1/2*-null *ad lib* | **-0.34  (-0.48, -0.17)** | **-11.6  (-14.7, -8.31)** | -0.31  (-0.92, 0.21) | -3.91  (-7.41, 0.87) |
|  | *Per1/2*-null TRF | 0.10  (-0.06, 0.23) | **-10.1  (-12.1, -7.21)** | -0.65  (-1.41, 0.92) | **-5.40  (-9.42, -2.17**) |
| **Expt. 2** | WT TRF  (reference group) | Reference group | Reference group | Reference group | Reference group |
|  | *Per1/2*-null *ad lib* | **-0.565  (-0.64, -0.50)** | **-5.26  (-6.84, -4.23**) | **-0.84  (-1.17, -0.53)** | **0.04  (-0.06, -0.01)** |
|  | *Per1/2*-null TRF | -0.065  (-0.14, -0.00) | -0.09  (-1.19, 0.99) | **-1.29  (-1.59, -0.91)** | **0.04  (-0.07, -0.01)** |

**Table S2**

**Mean effect sizes (with bias-corrected 95% confidence intervals) relative to control mice.** Mean effect sizes calculated using 5000 bootstrap resamples and are differences relative to ‘WT *ad lib*’ mice for Experiment 1 and ‘WT TRF’ mice for Experiment 2. In bold are effects in which the 95% confidence intervals do not include zero.

|  | Group | Mean starting  weight ±SEM  (g) | Mean ending  weight ±SEM  (g) | Mean  % weight loss | Mean difference in % weight loss  (95% CI) |
| --- | --- | --- | --- | --- | --- |
| **Expt. 1** | WT *ad lib* | 24.9 ± 0.6 | 24.0 ± 0.2 | 3.4 ± 1.8 | Reference group |
|  | WT TRF | 28.2 ± 0.5 | 25.6 ± 0.6 | 9.2 ± 2.2 | 5.9 (-0.9, 9.8) |
|  | *Per1/2*-null *ad lib* | 29.5 ± 1.0 | 26.0 ± 0.3 | 11.5 ± 3.2 | **8.1 (2.7, 15.6)** |
|  | *Per1/2*-null TRF | 24.8 ± 0.6 | 23.1 ± 0.4 | 6.7 ± 1.9 | 3.3 (-1.5, 7.5) |
| **Expt. 2** | WT TRF | 27.5 ± 0.6 | 25.5 ± 0.4 | 7.3 ± 1.3 | Reference group |
|  | *Per1/2*-null *ad lib* | 23.7 ± 0.7 | 21.5 ± 0.6 | 9.2 ± 2.4 | 1.9 (-1.7, 7.8) |
|  | *Per1/2*-null *TRF* | 23.5 ± 0.3 | 20.7 ± 0.4 | 12.0 ± 1.4 | **4.7 (1.6, 8.2)** |

**Table S3**

**Mean mouse weights and weight loss.** WT and *Per1/2*-null mice were given food either *ad lib* or had access to food restricted to 10 hours per day between 09:00 and 19:00 (time restricted diet, TRF). Starting measures were taken at Day 2 PI and ending weights at Day 6 PI. Mean effect sizes calculated using 5000 bootstrap resamples and are differences relative to reference groups: ‘WT *ad lib*’ mice for Experiment 1 and ‘WT TRF’ mice for Experiment 2. In bold are effects in which the 95% confidence intervals do not include zero.
